# Exploiting aberrant mRNA expression in autism for gene discovery and diagnosis

**DOI:** 10.1101/029488

**Authors:** Jinting Guan, Ence Yang, Jizhou Yang, Yong Zeng, Guoli Ji, James J. Cai

## Abstract

Autism spectrum disorder (ASD) is characterized by substantial phenotypic and genetic heterogeneity, which greatly complicates the identification of genetic factors that contribute to the disease. Study designs have mainly focused on group differences between cases and controls. The problem is that, by their nature, group difference-based methods (e.g., differential expression analysis) blur or collapse the heterogeneity within groups. By ignoring genes with variable within-group expression, an important axis of genetic heterogeneity contributing to expression variability among affected individuals has been overlooked. To this end, we develop a new gene expression analysis method—aberrant gene expression analysis, based on the multivariate distance commonly used for outlier detection. Our method detects the discrepancies in gene expression dispersion between groups and identifies genes with significantly different expression variability. Using this new method, we re-visited RNA sequencing data generated from post-mortem brain tissues of 47 ASD and 57 control samples. We identified 54 functional gene sets whose expression dispersion in ASD samples is more pronounced than that in controls, as well as 76 co-expression modules present in controls but absent in ASD samples due to ASD-specific aberrant gene expression. We also exploited aberrantly expressed genes as biomarkers for ASD diagnosis. With a whole blood expression data set, we identified three aberrantly expressed gene sets whose expression levels serve as discriminating variables achieving >70% classification accuracy. In summary, our method represents a novel discovery and diagnostic strategy for ASD. Our findings may help open an expression variability-centered research avenue for other genetically heterogeneous disorders.

## Introduction

Autism spectrum disorder (ASD, [OMIM 209850]) is a complex neurodevelopmental condition characterized by substantial phenotypic and genetic heterogeneity (Devlin and Scherer 2012; Geschwind 2011; Geschwind and State 2015; Willsey and State 2015). Both genetic and environmental factors contribute to the increased risk of developing ASD (Persico and Bourgeron 2006; Sandin et al. 2014). Recent years have seen major advances in the understanding of the genetic, neurobiological and developmental underpinnings of ASD (Abrahams and Geschwind 2008; Belmonte et al. 2004; Elsabbagh and Johnson 2010). Genetic studies, especially genome-wide association studies (GWAS), have identified many singlenucleotide variants (SNVs) and copy number variants (CNVs) associated with ASD susceptibility (Glessner et al. 2009; Wang et al. 2009; Weiss et al. 2009). However, it remains difficult to identify the actual causal genes underlying these associations. SNVs that produce association signals in identified loci often fall into intergenic regions, while CNVs often extend across multiple variants or genes, both of which confound the identification of causal genes. Also, there are opposing views on the relative contribution of rare versus common variants to ASD susceptibility. Some studies suggest that low-frequency variants bring a greater impact on the risk for ASD (Neale et al. 2012; Pinto et al. 2014; Sanders et al. 2012; Sebat et al. 2007), while other studies suggest that common variants form a dominating source of the risk (Gaugler et al. 2014; Klei et al. 2012). Against this background of complexity, several studies demonstrate the use of gene expression information—measuring mRNA abundance of individual genes, coupled with other genetic approaches, allows for novel insights in understanding ASD (Flint et al. 2014; Gupta et al. 2014; Voineagu et al. 2011). Analyzing gene expression and sequence data facilitates revealing the impact of regulatory genetic variants on the gene itself and the indirect consequences on the expression of other genes (Iossifov et al. 2014).

To this end, we introduce a novel, gene expression analysis method for identifying ASD-implicated genes. Our working hypothesis is that ASD is associated with aberrant gene expression caused by the promiscuous multigene activation and repression. Indeed, we show that many gene sets that contain genes known to be implicated in ASD tend to be expressed more aberrantly in ASD-affected individuals. Encouraged by these findings, we conduct a searching for unique combinations of genes for ASD diagnosis based on whole blood expression data. We use a greedy algorithm to solve the combinatorial problem of global search and identify three gene sets, each containing five genes, which can be used as classifier gene sets with high sensitivity and high specificity to discern gene expression specific to ASD patients. Altogether, our results refine the relationships between gene function and gene expression dispersion among individuals affected with ASD, providing new insights into the genetic, molecular mechanisms underlying the dysregulated gene expression in ASD. Our results also lay out the foundation for the utilization of gene expression dispersion as biomarkers for early diagnosis of ASD.

## Materials and Methods

### Gene expression data

Whole transcriptomes of 104 brain tissue samples (47 ASD and 57 controls) were previously determined using RNA sequencing by Gupta et al. (2014). The data had been deposited in the National Database for Autism Research (NDAR) under the accession code NDARCOL0002034. Among these samples, 62, 14 and 28 were tissues from cerebral cortex (Brodmann Area [BA] 19), anterior prefrontal cortex (BA 10), and a part of the frontal cortex (BA 44), respectively, resulting in 47 (32 unique individuals) ASD samples and 57 (40 unique individuals) controls. For this study, the raw data of gene expression was normalized using the conditional quantile normalization (Hansen et al. 2012) and then processed using the algorithm of probabilistic estimation of expression residuals (PEER) (Stegle et al. 2010) to remove technical variation. PEER residuals were obtained after regressing out covariates (age, gender, brain region, and sample collection site) and factors accounting for ten possible hidden determinants of expression variation. The expression median across all samples was added back to the PEER residuals to give the final processed gene expression levels. Extremely lowly expressed genes with expression median < 2.5 (empirical cutoff) were excluded. The final data matrix contained the expression level information of 10,127 genes in 104 samples. Principal component analysis was performed to indicate that there was no population stratification regarding the global gene expression profiles (**Supplementary Fig. 1**).

### Functional gene sets

The curated gene sets (n = 10,348) used in the Gene Set Enrichment Analysis (GSEA) were obtained from the molecular signatures database (MSigDB v5.0, accessed March 2015) (Liberzon et al. 2011). GO terms (n = 14,825) associated with protein-coding genes were downloaded from BioMart (v0.7, accessed February 2015) (Smedley et al. 2015). The coexpression networks were built for control samples using the Weighted Gene Co-expression Network Analysis (WGCNA) (Langfelder and Horvath 2008). The power of 16 was chosen using the scale-free topology criterion; the minimum module size was set to 4, and the minimum height for merging modules was 0.25. The resulting modules were plotted using SBEToolbox (Konganti et al. 2013). The list of ASD-implicated genes was obtained from the Simons Foundation Autism Research Initiative (SFARI) Gene Scoring Module. The list includes 410 genes in the categories S and 1 – 4, which stand for syndromic, high confidence, strong candidate, suggestive evidence, and minimal evidence, respectively, indicating the strength of the evidence linking genes to ASD. Of the 410 genes, 294 genes are in the expression data matrix we analyzed.

### Calculation of robust MD between ASD and control samples

For a given gene set, *MD_i_* is the Mahalanobis distance (Mahalanobis 1936) from an ASD individual *i* to the multivariate centroid of control individuals. Conventional *MD_i_* was calculated using the following operation:

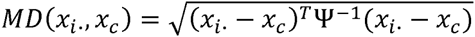

where *x_i_* is the vector of expression of genes in ASD sample *i, x_c_* is the vector of expression means of genes across all control samples, and ψ is the covariance matrix estimated from the controls. Throughout the study, a robust version of *MD_i_* was calculated using the algorithm Minimum Covariance Determinant (MCD) (Rousseeuw and Van Driessen 1999). The MCD algorithm subsamples *h* observations out of control individuals whose covariance matrix had the smallest covariance determinant. By default, *h* = 0.75*n*, where *n* is the total number of control samples. The robust *MD_i_* was then computed with the above equation by replacing *x_c_* with the MCD estimate of location, *x̂_c_*, i.e., the expression mean of the *h* controls, and replacing ψ with the MCD estimate of scattering, 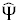, i.e., the covariance matrix estimated from the *h* controls. A Matlab implementation of MCD, available in the function `mcdcov` of LIBRA toolbox (Verboven and Hubert 2005), was used to perform the computation of MCD estimator. For a given gene set, MCD estimates of *x̂_c_* and 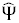 were computed as the outputs of `mcdcov`, and re-used for calculating robust *MD_i_* for ASD individuals.

The sum of squared 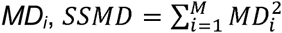 was calculated for given gene sets to measure the overall dispersion of *M* individuals affected with ASD. To assess the significance of SSMD of a given gene set, permutation tests were performed with *N* reconstructed gene sets of the same size but randomly selected genes. The *P*-value of permutation test, *P_perm_*, was determined by the ratio of 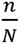, where *n* is the number of random gene sets having SSMD greater than that of the tested gene set and *N* is the total number of random gene sets used in permutation tests. To save computational time, we first set *N* = 1,000 to obtain a short list of gene sets with *P_perm_* < 0.001, and then set *N* = 10,000 to obtain nominal *P_perm_* for gene sets in the short list. The correction for multiple testing was performed by controlling the false discovery rate (FDR) with the Benjamini-Hochberg method (Benjamini and Hochberg 1995). To measure the relative contribution of each gene in a gene set to the total SSMD of the gene set, ΔSSMD was calculated. ΔSSMD is the difference between the two SSMD values calculated before and after the gene is excluded from the gene set.

### Receiver operator characteristic (ROC) curve analysis

For the analysis of aberrant gene expression as the biomarker for ASD diagnosis, peripheral blood gene expression data measured using the Affymetrix Human Gene 1.0 ST array for 104 ASD and 82 controls was downloaded (GEO accession: GSE18123) (Kong et al. 2012). The raw intensity data was processed using the R function `rma` (robust multi-array average expression measure) in Affy package. The expression measure was quantile normalized and log2 transformed. The final data matrix contained the expression level information for 16,365 autosomal genes among 186 samples. We equally split samples into a training set and a test set, each of which contains half of ASD (i.e., 52 ASD samples) and half of control samples (i.e., 41 controls).

ROC curve analysis was used to evaluate the specificity and sensitivity of classification tests, in which gene sets were used as classifiers for ASD and controls. For a given gene set, we first obtained the multivariate centroid of controls in the training set (***G***_*traimng*_) and calculated *MD_i_* of each sample *i* (including all ASD and control samples in the training set). Using ROC curve analysis, the threshold (denoted by *T*) corresponding to optimal specificity and sensitivity combination was determined. If *MD_i_* is greater than *T*, the sample *i* was classified as an ASD, otherwise a control. The performance of each gene set for predicting ASD and control samples was tested at different threshold *T* values to obtain the area under the curve (AUC). We denote the AUC values with respect to the training and test data sets as AUC1 and AUC2. After selecting three classifier gene sets with top AUC1 values, we then assessed their prediction performances with the test data set using ROC curve analysis again. For each classifier gene set, *MD_i_* for all samples of the test set, regardless of the disease status of samples, was calculated against ***G**_training_*, i.e., the multivariate centroid of controls in the training set.

### Global search for the classifier gene sets

A greedy algorithm was developed to identify subsets of genes, for which AUC1 reaches its maximal values as possible. The calculation of AUC1 can be seen in section “Receiver operator characteristic (ROC) curve analysis.” The search is global because combinations of all expressed genes were considered and no information of any pre-defined gene sets was used. Starting with all possible two-gene combinations, AUC1 values were computed, and the top 1,000 two-gene pairs with maximal AUC1 were retained as seeds for subsequent steps. The idea of the greedy strategy is to make a locally optimal choice at each stage have the hope of finding a global optimum. Thus, the assumption here is that the genes in the two-gene combinations producing the greatest AUC1 (i.e., the locally optimal solution) would be among those in five-gene combinations producing the greatest AUC1 (i.e., the global optimum). For each of the selected two-gene combinations, a new gene that can produce largest SSMD was identified and added to the gene pair to make a three-gene combination. The procedure was repeated until the number of genes reached five. At this stage, an additional procedure was introduced to improve the locally optimal solutions achieved by the greedy heuristic: all distinct three-gene combinations were extracted from five-gene combinations as the new candidate subsets. From these three-gene combinations, new genes were added to get a new round of solutions of five-gene combinations. The newly generated five-gene combinations will be retained if they produced larger AUC1 than older ones. This replacement procedure was repeated until no improvement could be made. For the top gene sets that produced the best AUC1 values, we then assessed their performances of prediction with the test data set. Computer code is available from the authors upon request.

## Results

Many biological processes underlying human diseases are accompanied by changes in gene expression in corresponding tissues (Cookson et al. 2009). ASD is not an exception. Previous studies have detected specific gene expression changes in ASD, concerning genes involved in the synaptic formation, transcriptional regulation, chromatin remodeling, or inflammation and immune response (Voineagu et al. 2011). These analyses mostly focused on departures across the average expression between the case and control groups, without considering or much less focusing on alternative patterns of departure such as those characterized by heterogeneous dispersion. The goal of present study is to detect the difference in heterogeneous multigene expression dispersion between samples derived from ASD-affected individuals and healthy controls.

### Overview of aberrant gene expression analysis

We have previously developed a multivariate method, namely aberrant gene expression analysis, to measure the level of multigene expression dispersion in the general population (Zeng et al. 2015). This analysis method uses Mahalanobis distance (MD) to quantify the dissimilarity in multigene expression patterns between individuals (Mahalanobis 1936). MD is an appropriate measure because it accounts for the covariance between expression levels of multiple genes. The aberrant gene expression analysis can be used to identify genes more likely to be aberrantly expressed among given individuals. It can also be used to identify individual outliers whose expression for a given gene set differs markedly from most of a population.

Here, we extend the MD-based aberrant gene expression analysis to a two-group setting. We estimated the level of gene expression dispersion among individuals affected with autism relative to controls, assuming that the increased dispersion is due to a promiscuous gene activation and repression associated with autism. We applied the aberrant gene expression with such a two-group setting and re-analyzed the gene expression data generated from the postmortem brain tissues of 47 ASD and 57 control samples (Gupta et al. 2014). For a given gene set, we computed the MD between gene expression of each ASD individual *i* to the multivariate centroid of the controls, denoted as *MD_i_*. We used the sum of squared *MD_i_* (SSMD) to measure the overall dispersion level for all ASD samples vs. the controls. Using permutation tests, we assessed the significance of gene sets and identified gene sets more likely to be aberrantly expressed among individuals affected with ASD (**Materials and Methods**).

### Coordinated gene expression is disrupted in ASD

Our MD-based aberrant gene expression analysis is capable of detecting the signal of expression aberration in different forms, including, e.g., (1) the increased individual-to-individual gene expression variance (i.e., the increased gene expression variability) and (2) the decreased expression correlation between genes. To illustrate the effect of disrupted expression, we use gene sets comprising only two genes to show the aberrant gene expression among individuals affected with ASD manifested as the loss of expression correlation between the two genes. **Fig. 1** shows scatter plots of expression levels between gene pairs. In **Fig. 1A**, the expression of *CORO1A* is positively correlated with that of *SYN2* for the controls (left panel, Pearson correlation test, *P* = 3.6×10^−10^). The gene *CORO1A* encodes coronin 1A, an actin binding protein. The gene *SYN2*, which is selectively expressed at nerve terminals in mature neurons, encodes synapsin II, a neuron-specific synaptic vesicle phosphoprotein (Cesca et al. 2010; Corradi et al. 2014). Synapsins interact with actin filaments in a phosphorylation-dependent manner (Benfenati et al. 1989). As evident from the description of gene functions, the correlated expression between the two genes is crucial for their respective actin-related molecular functions in normal individuals. However, such a crucial correlation becomes less significant in the ASD group (middle panel, *P* = 0.07). To make the contrast, we superimposed the data points for ASD individuals onto those of the controls (right panel). The top 10 ASD samples with the largest MD values are highlighted with red asterisks. The data points of these ASD samples are the most remote observations, distributed either far away from or orthogonally against the “cloud of data points” around the population mean, in which most control individuals are located. **Fig. 1B** presents a negative example, in which the correlations in expression levels between two genes, *CX3CR1* and *SELPLG*, are presented in both control and ASD groups (*P* = 1.3×10^−9^ and 1.4×10^−10^, respectively), indicating that the coordinated expression between the two genes is not disrupted in ASD. Altogether, these two examples, one positive and one negative, suggest that aberrant gene expression is not universal. The pattern of aberration may be highly specific with regard to certain gene sets (e.g., that in **Fig. 1A)** but not others (e.g., that in **Fig. 1B**).

**Fig. 1.**
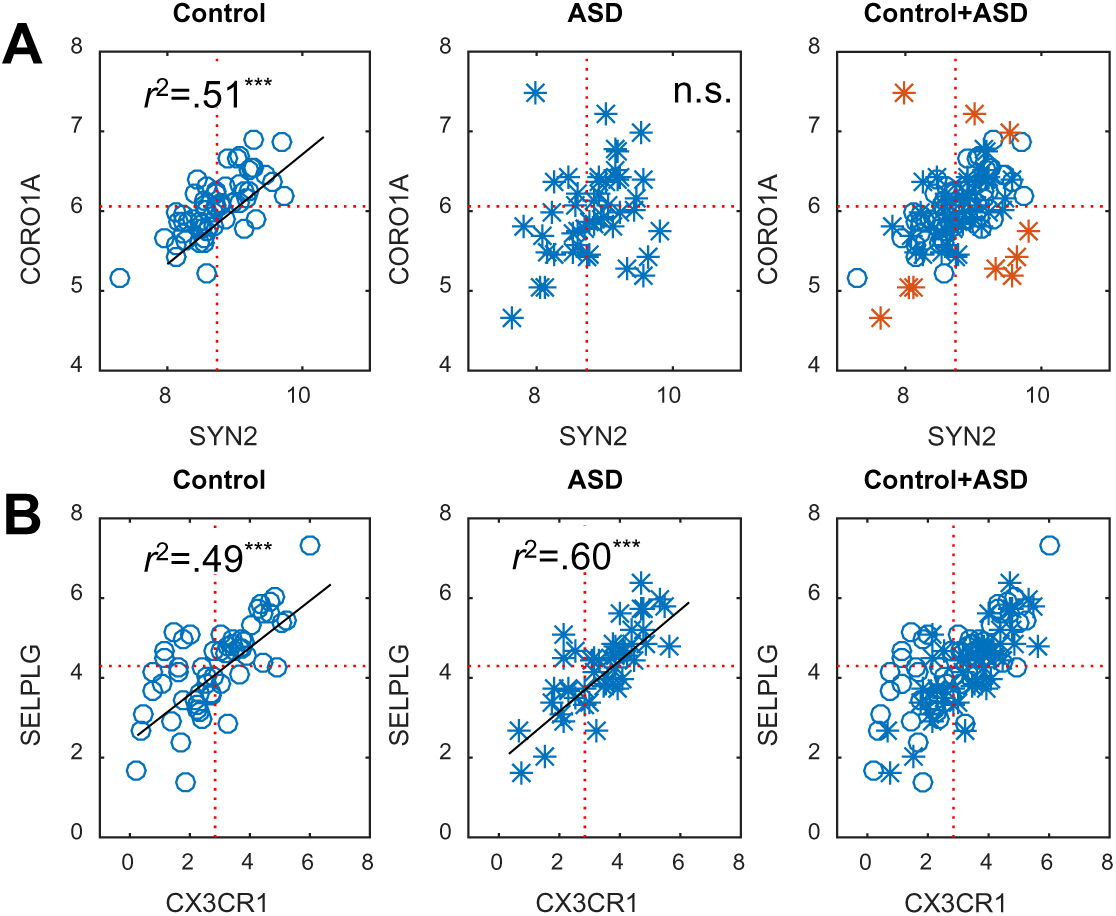
A proof-of-concept example, based on real data (Gupta et al. 2014), showing that (**A**) the correlated expression between *SYN2* and *CORO1A* presents among non-ASD samples but is disrupted among ASD samples, while (**B**) the correlated expression between *CX3CR1* and *SELPLG* presents among both non-ASD and ASD samples. Red stars in (**A**) show the top 10 ASD samples with the largest *MD_i_* and *r^2^* is the squared Pearson correlation coefficient.

### Functional gene sets that tend to be aberrantly expressed in ASD

To identify gene sets more likely to be aberrantly expressed in ASD samples, we calculated SSMD for a number of pre-defined gene sets (**Materials and Methods**). These included the curated gene sets in the MSigDB of GSEA (Liberzon et al. 2011). The significance of each gene set was assessed using permutation tests with random gene sets. A total of 18 GSEA curated gene sets were found to produce significantly higher SSMD than random gene sets at FDR of 10%. Functions of these gene sets fall into four major categories, namely, metabolism and biosynthesis, immune or inflammatory response, signaling pathway, and vitamins and supplements (**Table 1**). The relevance of these major functional categories with ASD is supported by respective studies (Abrahams and Geschwind 2008; Chow et al. 2012; Frye et al. 2010; Klaiman et al. 2013; Lazaro and Golshani 2015; Sawicka and Zukin 2012; Tierney et al. 2006). For example, the mTOR signaling pathway, which has a full name in the Reactome database: Energy dependent regulation of the serine/threonine protein kinase mTOR by LKB1-AMPK (Croft et al. 2014), is essential to synaptogenesis; gene products of the pathway regulate dendritic spine morphology in synapses. The dysregulation of this pathway is implicated in ASD (Abrahams and Geschwind 2008; Lazaro and Golshani 2015; Sawicka and Zukin 2012). **Table 1** also contains four gene sets with miscellaneous functions, unclassified into any of the four major categories but all implicated in ASD. These genes are involved in: (1) activated point mutants of *FGFR2* (Schubert et al. 2015; Stevens et al. 2010), (2) activation of the AP-1 family of transcription factors (Schaaf et al. 2011), (3) inwardly rectifying K* channels (Guglielmi et al. 2015; Lee et al. 2014), and (4) G2/M checkpoints (Fatemi et al. 2008).

**Table 1.** GSEA curated gene sets that tend to be aberrantly expressed in ASD. *Number of genes included in our analysis/Number of genes in the gene set. Gene sets mentioned in the main text are shown in italic. SFARI ASD-implicated genes are shown in bold.

| GSEA gene set | Number of genes* | Top ΔSSMD gene | Reference |
| --- | --- | --- | --- |
| Metabolism and biosynthesis |  |  |  |
| KEGG_PENTOSE_PHOSPHATE_PATHWAY | 19/27 | *H6PD, PRPS2, PFKP* |  |
| KEGG_STEROID_BIOSYNTHESIS | 14/17 | *SC5DL, NSDHL, **DHCR7*** |  |
| REACTOME_CHOLESTEROL_BIOSYNTHESIS | 20/24 | *SQLE, HSD17B7, HMGCR* | (Tierney et al. 2006) |
| REACTOME_BRANCHED_CHAIN_AMINO_ACID_CATABOLISM | 16/17 | *DLD, HIBADH, MCCC2* |  |
| Immune/Inflammatory response |  |  |  |
| BIOCARTA_LAIR_PATHWAY | 4/17 | *SELPLG, C3, ITGB1* |  |
| BIOCARTA_41BB_PATHWAY | 12/17 | *MAPK8, ATF2, MAPK14* |  |
| REACTOME_IL1_SIGNALING | 25/39 | *CHUK, RBX1, BTRC* | (Chow et al. 2012) |
| REACTOME_REGULATION_OF_IFNA_SIGNALING | 6/24 | *STAT1, PTPN1, JAK1* |  |
| Signaling pathway |  |  |  |
| BIOCARTA_IGF1_PATHWAY | 20/21 | *JUN, CSNK2A1, ELK1* |  |
| PID_S1P_S1P2_PATHWAY | 21/24 | *MAPK8, MAPK14, JUN* |  |
| PID_HNF3APATHWAY (FOXA1/HNF3A TF network) | 22/44 | *NDUFV3, PISD, FOS* |  |
| REACTOME_ENERGY_DEPENDENT_REGULATION_OF_MTOR_BY_LKB1_AMPK | 15/18 | *PRKAA1, CAB39, **TSC1*** | (Abrahams and Geschwind 2008; Lazaro and Golshani 2015; Sawicka and Zukin 2012) |
| Vitamins and supplements |  |  |  |
| BIOCARTA_VITCB_PATHWAY | 6/11 | *SLC2A3, COL4A2, SLC2A1* |  |
| REACTOME_TETRAHYDROBIOPTERIN_BH4_SYNTHESIS_RECYCLING_SALVAGE_AND_REGULATION | 9/13 | *GCHFR, PTS, AKT1* | (Frye et al. 2010; Klaiman et al. 2013) |
| Miscellaneous |  |  |  |
| REACTOME_ACTIVATED_POINT_MUTANTS_OF_FGFR2 | 4/16 | *FGF9, FGFR2, FGF1* | (Schubert et al. 2015; Stevens et al. 2010) |
| REACTOME_ACTIVATION_OF_THE_AP1_FAMILY_OF_TRANSCRIPTION_FACTORS | 10/10 | *MAPK14, **MAPK3**, ATF2* | (Schaaf et al. 2011) |
| REACTOME_INWARDLY_RECTIFYING_K_CHANNELS | 20/31 | ***KCNJ10**, KCNJ4, GNG4* | (Guglielmi et al. 2015; Lee et al. 2014) |
| REACTOME_G2_M_CHECKPOINTS | 22/45 | *MCM2, RFC5, RPA2* | (Fatemi et al. 2008) |

To determine individual gene’s contribution to the total SSMD of a gene set, we computed ΔSSMD for each gene. The ΔSSMD of a gene is the difference between SSMD values of a gene set before and after excluding the gene from the gene set. Top three genes with the largest ΔSSMD are given for all gene sets in **Table 1**. SFARI ASD-implicated genes are highlighted. To further investigate the relationship between gene function and aberrant gene expression, we grouped genes into gene sets, based on their cellular and molecular functions indicated by gene ontology (GO) terms associated with the gene function descriptions. A total of 36 significant GO terms at an FDR of 10% were identified (**Supplementary Table 1**). These terms are distributed in 22 biological processes (BP), 11 molecular functions (MF), and three cellular components (CC) sub-ontologies. The relevant processes include cellular response to stimulus, cellular metabolic process, cell morphogenesis and proliferation, regulation of intracellular transport and organelle organization, and tissue development. A close look at these significant GO terms revealed several that are implicated in ASD, e.g., *neuropeptide receptor activity* (GO: 0008188) (Ramanathan et al. 2004), *neuropeptîde binding* (GO: 0042923) (Baribeau and Anagnostou 2015; Lim et al. 2005), and *inhibitory synapse* (GO: 0060077) (Pettem et al. 2013; Tabuchi et al. 2007).

### Co-expression modules that tend to be aberrantly expressed in ASD

We also used the expression data from non-ASD controls to construct the co-expression networks. We customized the parameters of WGCNA (Langfelder and Horvath 2008) (**Materials and Methods**), instead of using the default values provided by the program, to construct as many as 807 network modules with relatively small size (4 to 110 genes). Such an adjustment of parameters was necessary because the core function `mcdcov` for MD calculation requires the size of gene sets (i.e., the size of modules) is no greater than the size of control samples (n = 57). Otherwise, the multivariate centroid of controls would not be able to be computed. For all modules containing 57 genes or fewer, we computed SSMD and used permutation tests to assess the significance of the modules. We identified 76 significant modules that tend to be aberrantly expressed in ASD samples (**Supplementary Table 2**). Many genes in these modules have functions in the central nervous system. For example, module 1 is enriched with genes closely associated with synapse and cell junction while module 5 is enriched with genes involved in regulation of neurogenesis/neuron differentiation. **Fig. 2A** shows the co-expression relationships between genes in modules one and five among control samples. The coexpression patterns in the two modules are absent in ASD samples (**Fig. 2B**). It is striking to observe such complete breakdowns of essential functional modules in ASD cases.

**Fig. 2.**
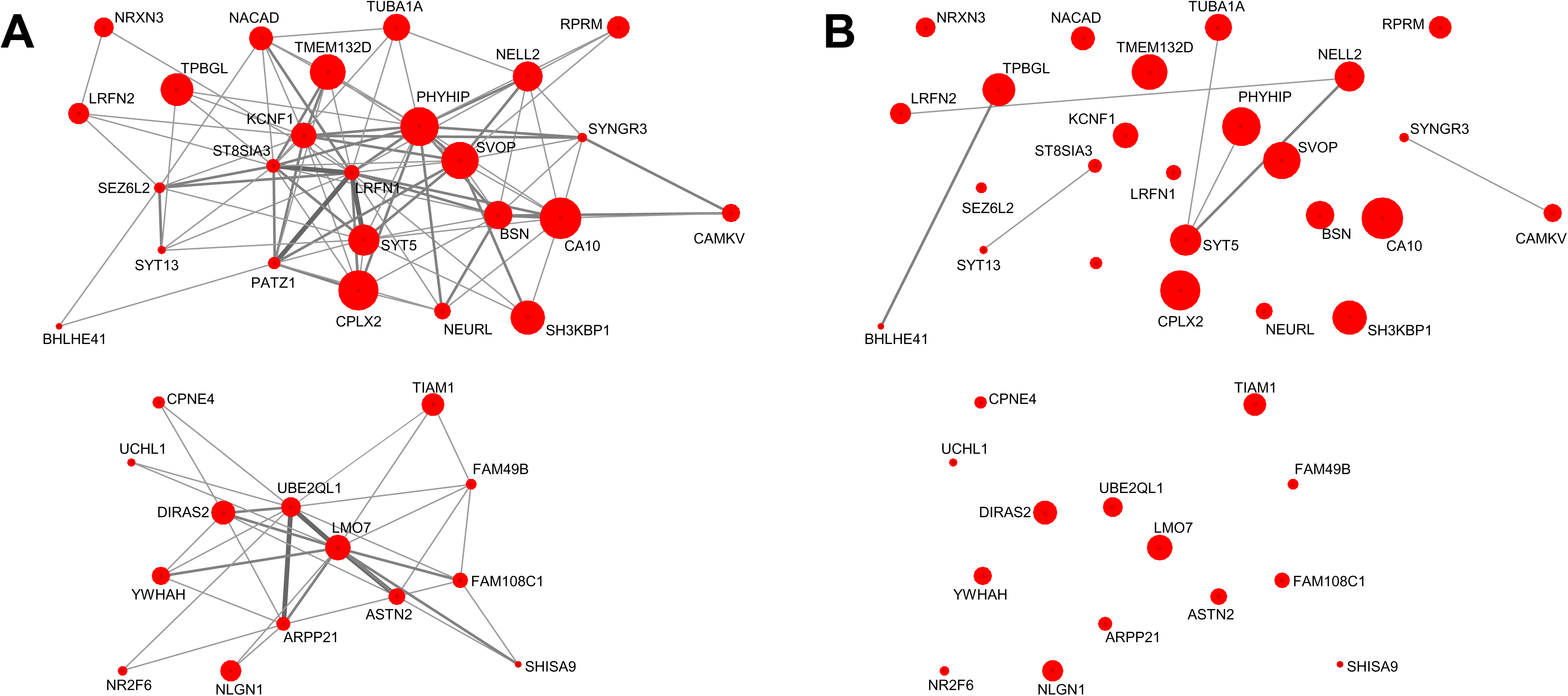
The breakdown of co-expression network modules in ASD. (**A**) Two example modules are presented as gene interaction subnetworks among non-ASD controls. Edge width is proportional to the value of Pearson’s correlation coefficient (ranging 0.5 – 0.8). Node size is proportional to the value of ΔSSMD for each gene. The two modules are enriched with genes whose products are closely associated with synapse or cell junction (top) and genes involved in regulation of neurogenesis or neuron differentiation (bottom), respectively. (**B**) The same sets of genes in the two modules are depicted for ASD samples. The missing of edges is due to the lack of co-expression relationships between genes.

To quantify the module difference between ASD and control groups, we used the function `modulePreservation` in the WGCNA R package (Langfelder et al. 2011) to calculate two statistics—medianRank and Zsummary—that measure the level of connectivity preservation between modules constructed using control and ASD samples. The majority of 76 significant modules have a large medianRank and a close-to-zero small Zsummary (**Supplementary Table 2**), which suggest little or no module preservation across the control and ASD samples. To demonstrate that the 76 significant modules constructed using control data are robust, we obtained an independent data from brain tissues of 93 non-ASD healthy controls (GEO accession: GSE30453) (Heinzen et al. 2008) and used this new independent expression data to re-draw these significant modules. We found that, despite the difference in technical platforms (i.e., RNA sequencing vs. microarray) on which two gene expression data were generated, most co-expression relationships between genes in these modules could be recapitulated using the new independent control data—representative modules that contain ten or more genes are shown in **Supplementary Fig. 2**. These results suggest that these modules are robust and the co-expression relationships between genes in these modules are biologically important and indispensable for healthy controls.

Next, we note that ΔSSMD may be used as a single-gene measure to prioritize genes with desired functions. For instance, *CPLX2* is among genes with the largest value of ΔSSMD in the module (**Fig. 2A**). It is likely that the sequences of *CPLX2* regulatory region are more heterogeneous among ASD samples, or the region contains variants associated with large gene expression variability more common in ASD samples. In either case, ΔSSMD enables to prioritize gene candidates and pinpoint the genomic regions that are likely to accommodate the potential mutations responsible for the increased gene expression variability. Indeed, *CPLX2* encodes a complexin protein that binds to synaphin as part of the SNAP receptor complex and disrupts it, allowing transmitter release. *CPLX2* has been associated with schizophrenia and attention deficit hyperactivity disorder (Lee et al. 2005; Lionel et al. 2011), but not with autism yet. In future, target sequencing of the *CPLX2* regulatory region in the ASD samples may allow us to discover unknown variants associated with autism risk. Alternatively, the deregulated *CPLX2* expression might be part of the dysregulation of the entire module, which could be due to a trans-regulatory change (e.g., a change in a regulator of *CPLX2* and associated module). In such a case, target sequencing may be used to rule out the influence of local regulatory mutations on the *CPLX2* expression.

We also tested the correlation between ΔSSMD and two network metrics for nodes, i.e., betweenness centrality and clustering coefficient. We previously showed that disease-causing genes have high betweenness centrality and low clustering coefficient values (Cai et al. 2010). However, for genes in these co-expression modules tested, no significant correlation was detected, which suggests ΔSSMD captured statistical features of genes that differ from those captured by the two network metrics. Finally, we examined whether, in the same modules, genes with large ΔSSMD tend to be expressed more differentially between ASD cases and controls. For genes in each of the 76 significant modules, we calculated t statistics using Student’s t-test to quantify the level of differential expression (DE) between ASD cases and controls. For each module, we then calculated the Spearman correlation coefficient (rho) between ΔSSMD scores and t statistics of all genes. The distribution of correlation coefficients for modules is symmetrical, centered at rho=0 with most values falling in between -0.5 and 0.5 (**Supplementary Fig. 3**), showing no consistent correlation between ΔSSMD scores and DE test statistics. Thus, DE is not predictive of ΔSSMD score or vice versa.

### Overlap between aberrantly expressed genes and ASD-implicated genes

Taking all pre-defined gene sets (i.e., GSEA-defined, GO term-defined, and WGCNA modules) together, a total of 10,127 genes were under the consideration of our gene set-based analyses, and 1,044 unique genes were present in the gene sets considered to be significant, for which gene expression profiles in ASD samples are over-dispersed. The overlap between these 1,044 genes and 294 SFARI ASD-implicated genes (**Materials and Methods**) is 36. These overlapping genes include eight of those belonging to the SFARI category of syndromic (*DHCR7, KCNJ10, MECP2, NF1, PAX6, SCN1A, TSC1* and *TSC2*), four in the category of high and strong confidence (*TBR1, ASXL3, BCL11A* and *DSCAM*), and 24 in categories of suggestive and minimal evidence. Two *de novo* loss-of-function mutations in *TBR1* have been previously identified in ASD patients (Neale et al. 2012; O'Roak et al. 2012a; O'Roak et al. 2012b), along with three in *ASXL3* (De Rubeis et al. 2014; Dinwiddie et al. 2013), two in *BCL11A* (De Rubeis et al. 2014; Iossifov et al. 2012), and four in *DSCAM* (De Rubeis et al. 2014; Iossifov et al. 2014). Nevertheless, the number of overlapping genes (36) is not significantly more than expected by chance (Hypergeometric test, *P* = 0.16). These results suggest that aberrant gene expression analysis, as a deviation from the *status quo*, produced the results of many novel candidate genes, which are not present in the gene list of current knowledge.

### Aberrant gene expression as biomarkers for ASD

We sought to determine whether we could classify patients as having ASD vs. controls solely based on the aberrant gene expression that is more pronounced in ASD samples. For this purpose, we obtained the gene expression data from the peripheral blood samples, including 104 ASD patients and 82 healthy controls (GEO accession: GSE18123) (Kong et al. 2012). The rationale of using this blood sample data set, rather than re-using the data set of post-mortem brains (Gupta et al. 2014), is from the position of the practical application. For diagnostic purposes, measuring gene expression in the peripheral blood makes more sense. Thus, in our analysis, a direct search for the biomarkers using the blood expression data is desired. After downloading the blood gene expression data (Kong et al. 2012), we split the data set into “training” and “test” sets, each containing data of 52 ASD and 41 control samples. With the training set data, we calculated *MD_i_* for ASD samples against the control samples. With the test set data, we calculated *MD_i_* for both ASD and control samples against the control samples of the training set (**Materials and Methods**). That is, we calculated *MD_i_* for all samples against the same set of controls in the training set.

Our purpose was to identify gene sets comprising several genes out of autosomal protein-coding genes expressed in the whole blood (*n* = 16,365) whose aberrant gene expression could be used to distinguish ASD cases from controls (i.e., *MD_i_* respecting the gene sets for ASD and non-ASD samples differs greatly). Here we used ROC curve analysis to evaluate the classification performance of a specific classifier gene set, so a search was conducted for gene sets with top AUC (the area under ROC curve) values based on the training set, for which the performances were assessed with the test set. We denote the AUC values with respect to the training and test data sets as AUC1 and AUC2. With randomly generated gene sets, we examined AUC2 as a function of the size of gene sets. We found no correlation between the two (**Supplementary Fig. 4**), suggesting that random gene sets have no prediction value. The positive correlation between the size of gene sets and AUC1 (**Supplementary Fig. 4**) is simply because that the inclusion of more variables (i.e., expression data from more genes) allows a better fit to the data. That is, the expression variation in control samples of the training set is better explained by more genes to be considered, resulting in a continuously improved AUC1. Nevertheless, the overfit of data for the training set (better AUC1) did not necessarily contribute much to the prediction for the test set (better AUC2), as suggested by the weak positive correlation between AUC1 and AUC2 (**Supplementary Fig. 5**). Based on these preliminary results, we decided to search for gene sets containing as few as five genes to avoid the potential problem associated with overfitting of the control data. Computational time is another consideration—an exact solution for such a search for more than five genes is a combinatorial problem requires >10^18^ SSMD calculations, which is computationally infeasible.

To search for the five genes, we developed a greedy algorithm to search from different starting points for producing local optimal solutions (**Materials and Methods**). Three gene sets, each containing five genes, were identified to generate high accuracy with balanced sensitivity and specificity values for the tests using both training and test data sets (**Fig. 3**). These gene sets are: {*FAM120A, HDC, OR13C8, PSAP, RFX8* }, {*HBG1, MOCS3, PDGFA, SERAC1, SLFN12L* }, and {*BHMT2, CCL4L1, CD2, FAM189B, MAK* } (see **Supplementary Table 3** for corresponding SSMD and ΔSSMD values). All three gene sets achieved greater than 70% sensitivity and greater than 70% specificity in all tests (**Table 2**). Further analyses showed that the prediction power of the three gene sets largely remained no matter how the original data (Kong et al. 2012) was randomly split into training and test sets. Some of these classifier genes are associated with ASD, although mostly in an indirect manner. For instance, the protein product of *FAM120A* interacts with that of the ASD-implicated gene *CYFIP1* (De Rubeis et al. 2013). A rare functional mutation in *HDC*, which encodes L-histidine decarboxylase catalyzing the biosynthesis of histamine from histidine, has been associated with Tourette’s syndrome (Ercan-Sencicek et al. 2010)—a neuropsychiatric disorder potentially related to ASD (Clarke et al. 2012). The expression of *PDGFA* was found to be down-regulated in patients affected with the 22q11.2 deletion syndrome, which is associated with high rates of ASD in childhood (Jalbrzikowski et al. 2015).

**Fig. 3.**
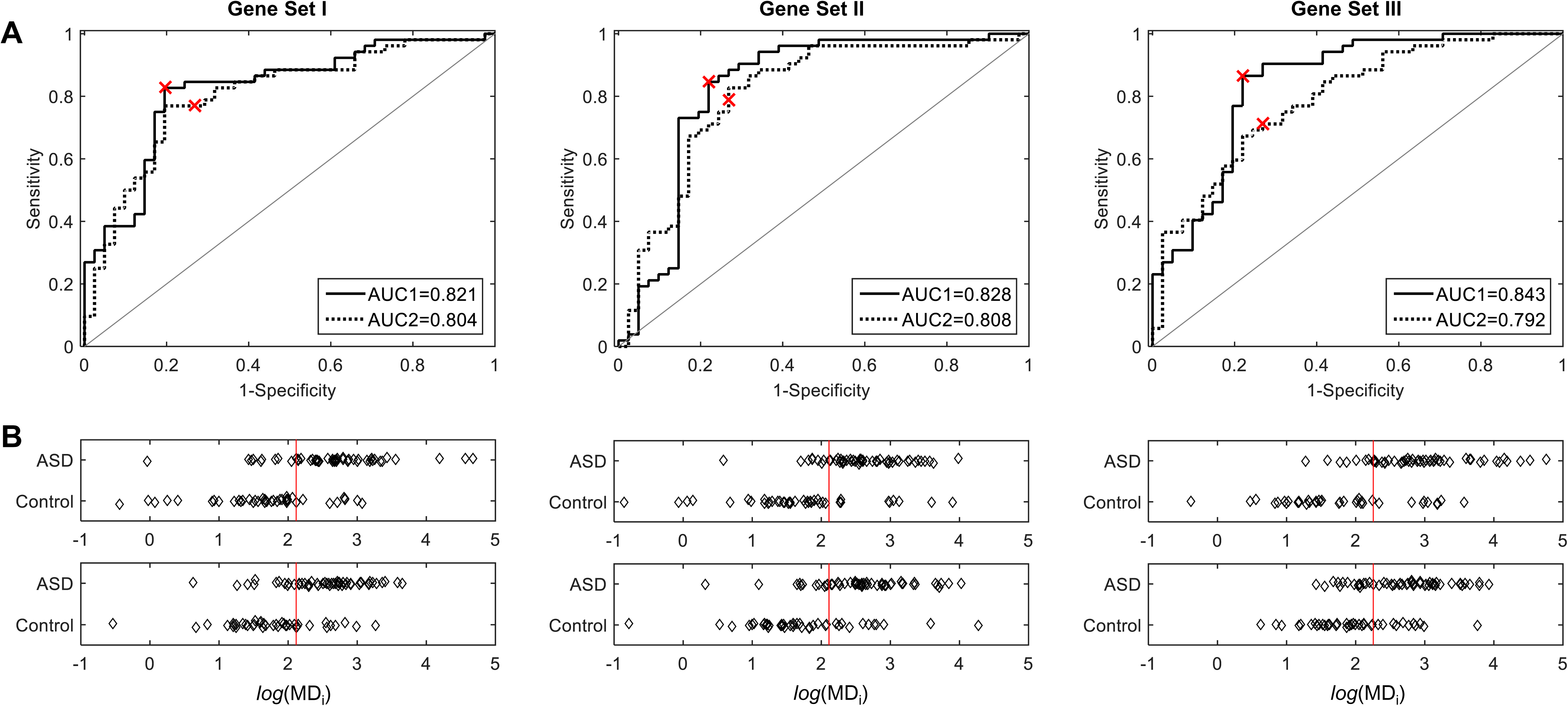
ROC curves and dot diagrams of *MD_i_*. (**A**) ROC curves graphs for the three classifier gene sets tested with the training and test data sets. Corresponding AUC values for the training (AUC1) and test (AUC2) data sets are given in the inserts. Red cross indicates the optimal operating point of the ROC curve for the training data set. (**B**) Dot diagrams for training (top) and test (bottom) sets showing the distributions of *MD_i_* calculated with respect to the three classifier gene sets for samples in ASD and control groups. Log-transformed *MD_i_* values are shown. The red vertical lines show the optimal cutoff values determined from the ROC curves tested on training data set.

**Table 2.**
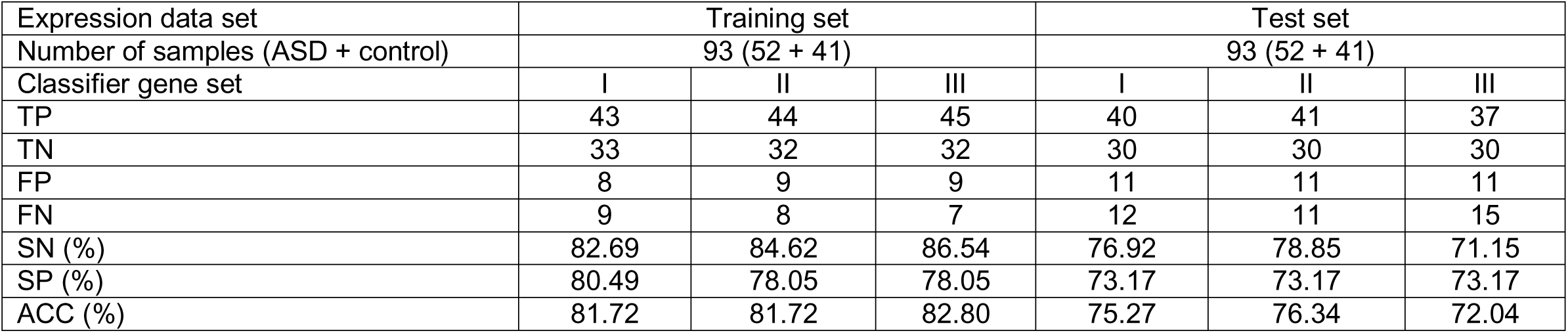
The performances of classifiers based on gene set I, II, III tested on the training and test data sets. True positive (TP), true negative (TN), false positive (FP), false negative (FN), sensitivity (SN), specificity (SP), accuracy (ACC) values are reported.

Finally, we repeated the searching for classifier gene sets using the expression data of brain samples (Gupta et al. 2014). Three classifier gene sets were obtained: {*IFI6, MIDN, MAPK8, ENO2, GYS1* }, {*HSPH1, ASH1L, IFIT3, GPR3, PCSK2* }, and {*HNRNPK, GOLT1B, BAZ2A, TRABD2A, UNG* }, which all gave ~75% or better prediction accuracy (**Supplementary Table 4**). These classifier genes show no overlap with those derived from the blood samples, but several are directly associated with ASD. For example, ASD-associated mutations have been identified in *ASH1L*, which is a SFARI category 1 (high confidence) gene (De Rubeis et al. 2014; Iossifov et al. 2014; Tammimies et al. 2015; Willsey et al. 2013) and in *MIDN*, which is involved in neurogenesis and neuronal migration (Butler et al. 2015).

## Discussion

ASD is a complex disease involving multiple genetic alterations that result in modifications of many cellular processes. Maladaptive patterns of ASD lead to significantly high gene expression variability among affected individuals. Unitary models of autism brain dysfunction have not adequately addressed conflicting evidence, and efforts to find a single unifying brain dysfunction have led the field away from research to explore individual variation and micro-subgroups. Therefore, it has been suggested that researchers must explore individual variation in brain measures within autism (Geschwind 2008; Waterhouse and Gillberg 2014). Previous studies, with few exceptions (Garbett et al. 2008; Voineagu et al. 2011), have rarely addressed the issue of increased gene expression variability associated with autism. Noticeably, Voineagu et al. (2011) pointed out: “Autistic subjects display significant phenotypic variability which could be due to an intricate interplay of genetic and environmental factors. Thus, we hypothesized that this phenotypic diversity is due to subject-to-subject variability in gene expression.” Nevertheless, the *status quo* pertaining to gene expression specific to ASD patients is based on the detection of differential gene expression, i.e., the gene-expression differences between mathematical expectation (i.e., mean) of ASD and control samples. The major assumption underlying differential expression analysis is: ASD cases have the same or similar gene expression change phenotypes, which makes them as a separate cohort have significantly higher or lower expression than the controls. However, this assumption contradicts the fact that ASD has highly heterogeneous genetic causes, and excludes empirical evidence gathered about uncommon molecular changes causing ASD (Neale et al. 2012; Pinto et al. 2014; Sanders et al. 2012; Sebat et al. 2007).

### Dispersion-specific measure of gene expression for autism

Our overall strategy for this study was based on the quantitative measures of the departure of multigene expression dispersion between individuals. The profound heterogeneity in ASD underscores the importance of leveraging measures of dispersion in order to capture the specific tendency. Gene expression dispersion has been found associated with gene function and disease or physiological status of individuals (Ecker et al. 2015; Li et al. 2010; Mar et al. 2011; Somel et al. 2006). Discrepancies in gene expression should not only be characterized by the mean but also by other statistics of interest, such as dispersion parameters. Using the proven multivariate approach (Zeng et al. 2015), we have further developed MD-based aberrant gene expression analysis and applied it to ASD. The statistical signal captured is the tendency of being more dispersed in multigene expression among ASD than control samples. We have shown that our variability-centered method can recapitulate signals from many genes known to be implicated in ASD. Our method does not depend on the prior knowledge about gene function or the identification of mutations in genes. Thus, it is a tool for discovering and identifying genes previously unknown to be involved in ASD progression.

### Aberrant gene expression in co-expression network modules

We have shown that, when applied to the co-expression network, SSMD can reveal the effects of perturbing genetic networks. SSMD analysis informs us about how ASD distorts expression patterns of biological systems. Disturbed ASD genetic networks have been noticed previously (Hormozdiari et al. 2015; Li et al. 2014; Parikshak et al. 2013; Pramparo et al. 2015; Willsey et al. 2013). However, most existing network analyses were not designed for directly measuring the level of dysregulation. Instead, information about known ASD genes, e.g., in (Li et al. 2014; Willsey et al. 2013), or differently expressed genes, e.g., in (Pramparo et al. 2015), were used to prioritize the modules, which would not allow modules contain unknown ASD genes to be prioritized for subsequent analyses. In contrast, our approach allows for a straightforward screening of perturbed network modules and provides the raw material for the identification of genetic regulatory mechanisms involved in the variability of gene transcription.

### Toward the genetic basis of aberrant gene expression

Our results have provided unique entry points to investigate further on the genetic basis of aberrant gene expression (e.g., increased gene expression variability) in ASD. When genotype or sequence information, along with their gene expression information, become available for ASD samples, it would be possible to assess the influences of the aggregation of rare mutations, CNVs, as well as common genetic variants on aberrant patterns of gene expression in ASD. In line with this view, the latest genome sequencing effort for autism-affected families showed that disruptive de novo/private mutations and CNVs are significantly enriched in regulatory regions of ASD-related genes in ASD probands (Turner et al. 2016). Furthermore, we have shown previously that certain common genetic variants, in addition to rare variants, cause the increase of gene expression variability among individuals (Hulse and Cai 2013). These common variants influence the variability of gene expression through the action of either epistasis or direct destabilization (Wang et al. 2014). By taking both rare and common variants into account, it would be possible to superimpose their impact onto a gene expression variability network to predict which parts of the network are more vulnerable to the perturbation from genetic factors such as ASD-related disruptive mutations.

### Aberrant gene expression as biomarkers

Recent years have seen an intensive search for biological markers for ASD. Although a wide range of ASD biomarkers has been proposed, as of yet none has been validated for clinical use (Walsh et al. 2011). Therefore, there is a critical need for valid biological markers for ASD.

Based on the results of aberrant gene expression analysis shown here, gene sets with just a few selected genes can be used as novel biomarkers. Application of our gene-expression candidate biomarkers will allow for higher sensitivity and specificity in a diagnostic screen for ASD. We anticipate that if our gene-expression biomarkers are expanded to use the blood gene expression data derived from other platforms (such as different types of microarrays, RNA sequencing, and qPCR), they will offer a significant advancement in developing a clinical blood test. The success of such gene-expression biomarkers will assist in early and objective diagnosis for ASD.

### Caveats and future directions

Voineagu et al. (2011) showed that the heterogeneity in gene expression between different brain regions of the same individual might introduce another level of gene expression variability. Due to the limitation of tissue samples available for this study, such an effect was not explicitly captured by our aberrant gene expression analysis. Also, brain regions themselves are highly heterogeneous because of the mixtures of cell types. Aberrant gene expression patterns might in part indicate different relative proportions of cell types in a sample. With the advent of the single-cell based technologies (Dey et al. 2015), this level of gene expression heterogeneity may be measured. Thus, the problem of heterogeneity of cell types in tissue samples as an important source of variability may be addressed in future studies.

### Conclusions

We have developed a novel, variability-centric gene expression analysis, and applied the method to ASD. This advance showcases the value of development and refinement of systems genomics tools in studying human complex diseases. The aberrantly expressed genes identified in this study will facilitate the identification of ASD-predisposing variation, which may eventually reveal the causes of ASD and enable earlier and more targeted methods for diagnosis and intervention.

## Authors’ Contributions

JJC conceived and designed the study. JG, EY, JY and ZY conducted the analyses. JG, EY, GJ and JJC wrote the manuscript.

## Supplementary Files

**Supplementary Table 1.** GO term-defined gene sets that tend to be aberrantly expressed in brain tissues of ASD-affected individuals. Gene sets contain genes annotated with GO terms of three sub-ontologies: biological process (BP), molecular function (MF), and cellular component (CC).

**Supplementary Table 2.** WGCNA co-expression network modules containing genes that tend to be aberrantly expressed in the brains of ASD-affected individuals. Modules are annotated with the DAVID-defined gene function keyword clusters. Representative genes with the corresponding function are shown in bold. Statistics of the preservation between modules built for cases and controls, medianRank and Zsummary, calculated using function `modulePreservation` of WGCNA are given.

**Supplementary Table 3.** Genes in the three classifier gene sets obtained from blood data set (GEO accession: GSE18123) and corresponding SSMD and ΔSSMD values.

**Supplementary Table 4.** Genes in the three classifier gene sets obtained from brain data set (57 controls and 47 ASD cases) and corresponding SSMD and ΔSSMD values. The performances of classifiers based on gene set I, II, III tested on the training and test sets are also reported including sensitivity (SN), specificity (SP) and accuracy (ACC) values.

**Supplementary Fig. 1.** Results of principal component analysis (PCA) showing the first four principal components (from PC1 to PC4). The distributions of 104 samples (57 controls and 47 ASD samples) on PCA spaces defined by PC1 and 2, PC2 and 3, and PC3 and 4 are shown.

**Supplementary Fig. 2.** Reproducibility of co-expression modules in the non-ASD control group and the breakdown of modules in ASD. Ten example modules are shown with two independent data sets from controls, as well as one data set from ASD samples. Edge width is proportional to the Pearson’s correlation coefficients (ranging 0.5 and 1). Node size is proportional to ΔSSMD for each gene.

**Supplementary Fig. 3.** Distribution of correlation coefficients between t statistics of DE test and ΔSSMD values of genes in 76 significant modules. The kernel density estimate of the distribution is shown with the gray line; values of Spearman correlation coefficient (rho) of modules are shown with orange triangles; rho=0 is shown with the dotted vertical line.

**Supplementary Fig. 4.** Box plot of AUC (area under ROC curve) value against the size of classifier gene set. For each size of the gene set (from 3 to 15), 100 different random gene sets were constructed and tested on the training set and test set for obtaining AUCs. The black and red boxplots denote AUC values tested on the training set (AUC1) and test set (AUC2) varying with the size of classifier gene set, respectively.

**Supplementary Fig. 5.** Scatter plot of AUC values tested on the test set (AUC2) against AUC values tested on the training set (AUC1) for 100 different random classifier 5-gene sets. Red line denotes the least-squares line of the scatter plot. The Spearman correlation coefficient between AUC1 and AUC2 is 0.32 (*P* = 1.1×10^−3^). The inset shows the distribution of the Spearman rank correlation coefficients between AUC1 and AUC2 calculated with 1,000 replicates of such 100 random classifier 5-gene sets.

## Acknowledgements

We thank Shannon Ellis and Dan Arking for sharing the data, Oliver Stegle and Tuuli Lappalainen for helping with data normalization, and Steve Horvath for the co-expression network analysis. We thank Rae L. Russell for proofreading and editing this paper. We acknowledge the Texas A&M Institute for Genome Sciences and Society (TIGSS) for providing computing resources and system administration support. This work was supported by the fund of China Scholarship Council to JG, and the National Natural Science Foundation of China (No. 61573296), the Specialized Research Fund for the Doctoral Program of Higher Education of China (No. 20130121130004), the Fundamental Research Funds for the Central Universities in China (Xiamen University: Nos. 201412G009, 201510384106) to GJ.

